## Supplementary material for "Neuron tracing and quantitative analyses of dendritic architecture reveal symmetrical three-way-junctions and phenotypes of *git-1* in *C. elegans*": SI text

### Software

The software was developed in MATLAB R2020b.

Toolboxes used: Navigation Toolbox, Robotics System Toolbox, Image Processing Toolbox, Curve Fitting Toolbox, Signal Processing Toolbox, Statistics and Machine Learning Toolbox.

The software and source code are available on GitHub:

https://github.com/Omer1Yuval1/Neuronalyzer

The GitHub commit id of the software version used to analyze the data and generate the figures is 5b62201 (March 03, 2021).

### Summary of the pipeline

Maximum intensity z-projection of the stacks are created for each neuron image. Images are rotated (and, if necessary, inverted) using Fiji (imageJ, NIH) such that the worm's head points to the left, and the ventral side points down.

Each image is then fed into a pre-trained convolutional neural network (CNN) designed for semantic segmentation, resulting in a binary image classified into neuron and non-neuron pixels (Fig. S1). The advantage of using a neural network stems from its ability to use intensity, shape and spatial context in order to detect morphological features, compared to classic morphological operations and filtering methods that are local and only consider pixel intensity and/or manually predefined shapes. Apart from the background, the CNN successfully distinguishes neuronal elements from other objects with similar intensities, such as gut granules (Fig. 2A). The CNN-derived binary image can be manually corrected for false positives and true negatives, although this step is not necessary (Fig. S2B). The binary image is then skeletonized using build-in MATLAB functions. The skeleton is used to find the approximated positions of neuronal vertices (junctions and tips), and then the binary image is used for finding their precise position and geometry. Next, the neuron is traced and reconstructed by fitting rectangular elements along its dendritic processes in the raw grayscale image. Here, the vertex positions are used as starting points and the CNN-derived skeleton is used as a constraint for the orientation of the fitted elements, in order to avoid false positives (Fig. 2B) and ensure that only neuronal elements are traced. Finally, the reconstructed neurons are saved into an abstracted database, used to extract morphological features and compared statistically across groups of animals.

**Machine learning**

Convolutional neural networks (CNN) are commonly used for computer vision problems, as they use spatial information to process an input. This is done by computing the convolution of filters over an image, and propagating this information through a series of hidden layers while spatial information is preserved. The network gets pairs of raw and annotated images, and optimizes its parameters to best predict the annotation based given the raw data [1,2].

We trained a convolutional neural network (CNN) for image segmentation based on SegNet (Fig. S1) [3]. The SegNet architecture was specifically designed for semantic segmentation of images, and its output is an image that matches the size of the input image where pixels are classified into one of predefined classes. We defined two pixel classes: neuron and non-neuron, which makes the output equivalent to a binary image. The size of the input, output and network parameters were adjusted to our custom problem (Table S1). For training, we used 3 annotated images of complete wild-type PVD of one-day adult worms. For each image, 5000 64x64 pixel image patches were chosen randomly, with a lower probability for samples that do not contain neuron pixels. Random reflections and rotations (in multiples of 90°) were applied to each sample (without duplications – each region was used only once, although patches are allowed to partially overlap). 80% of these were used for training and 20% for validation. The trained network performed with 98.07% accuracy on the validation set, compared to 98.37% on the training set. Following training, the CNN was applied to our dataset of 10 wild-type and 10 mutant PVD images, as well as to 10 additional images of wild-type PVDs of L4 and young-adult worms. The images that were used for training were not included in our datasets for morphological analysis. For each PVD image, the CNN was applied to unique 64x64 pixel image patches, using the "semanticseg" MATLAB function. This resulted in a binary image the same size as the raw PVD image, classified into neuron and non-neuron pixels (Fig. 2A and S2B).

Since all images were acquired using the same magnification, no further scaling is required. The size of the CNN-input image patch was chosen such that it is as small as possible to allow for a high number of distinct samples, yet contains complex neuronal structures in order for the CNN to have enough context to learn from.

Network architecture

The SegNet architecture is made of an encoder and decoder subunits that perform the convolution of the input, followed by a softmax layer that assigns class probabilities, and a pixel classification layer that computes the loss during training and performs the classification of new data [3].

The input layer gets small patches of grayscale neuronal images of a fixed size (here 64x64 pixels) and applies zero-centering normalization to them. The input is then processed through the encoder subunit that consists of a sequence of convolution units (Fig. S1, blue squares), followed by a max pooling layer (Fig. S1, green square). This sequence can be repeated, but is limited by the size of the input image (here we used a depth of 3). In the decoder subunit, the process is revered, with an upsampling layer (Fig. S1, purple square) replacing the pooling layer and preceding each set of convolution units.

Each convolution unit (blue square) consists of a 2D convolution layer, a batch normalization (BN) layer and a rectified linear unit (ReLU). The convolutional layers perform the convolution of a set of preinitialized filters of a fixed size (here 3x3 pixels) over an input matrix. The result is then normalized across the entire mini-batch by the BN layer, and then fed into the ReLU layer that sets all negative weights to zero. Finally, the pooling layer takes the maximum value of each 2x2 subregion of its input, resulting in a matrix that is four times smaller than the input matrix. The upsampling layer increases the size of its input by performing deconvolution of the corresponding layer in the encoder (Fig. S1, blue lines).

Finally, the softmax layer (Fig. S1, yellow square) takes the output of the last ReLU layer and computes its probability for belonging to each class c:

$$\sigma_c(\vec{z}) = \frac{e^{z_c}}{\sum_{i=1}^{C} e^{z_i}}$$

Where z is the output from last ReLU layer, and C is the number of classes. The result is a probability distribution that sums up to 1.

The last layer of the network is a pixel classification layer (Fig. S1, red square). It converts the output of the softmax layer into one of the predefined classes. During training, this layer computes the loss between the input and output for k mutually exclusive classes using the cross-entropy loss function.

**Detection of vertices (neuronal junctions and tips)**

Following image classification by the CNN, it was skeletonized using built-in MATLAB functions (Fig. S2C). In the skeleton binary image the PVD is reduced to a 1-pixel thick structure. The skeleton is further processed and pruned to remove processes and loops that are below detection limit and that do not correspond to neuronal structures.

The skeleton is first used to find the approximate positions of neuronal vertices. This is done using the build-in MATLAB function “bwmorph" with input parameters’ branchpoints’ and ’endpoints’ (corresponding to junctions and tips, respectively). This operation finds pixels that are connected to either 3+ pixels and 1 pixel (respectively) in their 8-connected neighborhood.

This information is then used, together with the binary image, to find the precise vertex positions and their geometry, defined using the vertex center point, radius and angles of its segments (Fig. 2J-M). The precise center of each vertex is detected by generating a dense 2D mesh around the approximated center, and convolving a growing concentric circle around each, against the binary image. The center point with the largest concentric circle that fully lies on neuron pixels (value = 1) is chosen as the corrected center point, and its radius is assigned as the vertex radius (Fig. 2J). To find the angles of the vertex, a rectangle is then convolved against the binary image by sliding it along the circumference of the vertex such that it points out (Fig. 2K). The peaks in this convolution function are then matched pairwise with the reference skeleton orientations, and used to define the angles of the vertex (Fig. 2L-M). Finally, the width of each rectangle is computed.

### Neuron tracing

To trace the processes of a neuron, it is first broken down into segments, defined as neuronal processes connecting two vertices. The vertices found in the previous step are used as starting points for the tracing of each segment. The skeleton image is used as a constraint to ensure that the path of each traced segment in the raw image is close to its corresponding skeleton segment.

First, each segment is matched with two vertex-derived rectangles. These rectangles define the initial tracing position and orientation, as well as the initial local width of the fitted rectangular element. Next, each of the starting rectangles is convolved against the grayscale image by rotating it around its origin (Fig. 2C-D). This convolution function is then normalized to the local background, sampled using two rectangles parallel to the neuronal process on both sides of the rectangle found in the previous step (Fig. 2G-H). This allows to use global parameters for peak analysis in the convolution function. Since multiple peaks may be detected, but only one is required, the peaks are sorted according to the resulting proximity to the specific skeleton segment. This allows to filter out peaks that capture non-neuronal elements or neighboring segments, and to choose the orientation that most likely corresponds to the segment's path. If no peaks are detected, or they do not meet the skeleton constraint, then the tracing of the segment is terminated. This helps avoid false positive segment paths. Finally, the width of the rectangle is adjusted to the local apparent width of the neuron (Fig. 2E-F). This is important to find the precise orientation in the next step when fitting the rectangular element to the neuronal process. The length of the convolution rectangle is a predefined function of its width (set globally as a length-width ratio parameter). This procedure is then repeated sequentially by incrementing the rectangle's origin by a fixed step size in the direction of the convolution-derived angle, until the two paths meet (Fig. 2B). Early termination of the tracing of a segment is completed using the skeleton.

**Simulation and fit of junction angle distribution**

We model PVD junctions as having an intrinsic geometry, characterized by the three junction angles $(\alpha_1, \alpha_2, \alpha_3)$. We further model the flexibility of the junctions by adding a gaussian noise term, $\sigma_\alpha$, to the orientation of the processes emanating from the junction. Each possible junction geometry, $J_i$, is therefore characterized by the ordered triplet of junction angles and the noise, $J_i = \left(\alpha_1^{(i)}, \alpha_2^{(i)}, \alpha_3^{(i)}, \sigma_\alpha^{(i)}\right)$. For every combination $J_i$ we generate $N$ junction realizations by simulating a gaussian random noise with a standard deviation $\sigma_a^{(i)}$. Importantly, while the random deviations of the processes emanting from the junction are normally distributed, the resulting distribution of junction angles is not so. We calculate the cumulative distributions of the three simulated junction angles, $F_{\alpha_1}(J_i), F_{\alpha_2}(J_i), F_{\alpha_3}(J_i)$. Finally, we compare the cumulative distributions of the simulated junction angles with the cumulative distributions of the angles extracted from the PVD images, $F_{\alpha_1}^{PVD}, F_{\alpha_2}^{PVD}, F_{\alpha_3}^{PVD}$ using a two-sample Kolmogorov-Smirnov test (Matlab ktest2.m), with the residual error defined as the sum of squares of the test statistics. The residual error phase plot was subsequently smoothed by Lowess fitting. The combination $J_i$ that minimizes the residual error is chosen as the best fit for the microscopy data. As a complementary analysis to the distribution of orientation of rectangular elements fitted to neuronal processes (θ), we show the same distribution for junctional elements (Fig. S3C). The differences between these two distributions (and in particular the peak at 13.5° for junctions, compared to the peak at 0° for dendritic processes) further strengthen our results that suggest that the orthogonal organization of dendritic processes does not arise from boundary conditions applied at the junctions (Fig. 5 and 5S).

### Visualization and statistical analysis

The software includes a user-interface and provides various options to visualize and validate the resulting neuron reconstruction and extracted morphological features.

The software also provides various interactive plots. These can be used to quantify morphological features of single and multiple neurons and compare them across datasets. This includes neuronal length, junction and tip count and density, vertices angles, curvature, midline orientation and azimuthal position. Interactive data manipulation options allow the user to break down quantities into their morphological classes, apply different normalizations, and adjust plot parameters such as histogram bin size. Finally, both the code and database are simple to access and interpret, making it easy to add custom quantifications and plots for the analysis of PVD or other neurons. A detailed user manual is available on the main page of the GitHub repository: https://github.com/Omer1Yuval1/Neuronalyzer.

### Curvature

The line curvature of a dendritic process is estimated from the smoothed discretized representation of rectangular elements $\{\vec{v}_i\}$ [4]. Boundary elements (belonging to junctions and tips) are excluded.

The curvature, $c_i$ , is estimated as the inverse of the circumscribed circle radius at a location $\vec{v}_i$ using the neighboring coordinates $\vec{v}_{i-1}, \vec{v}_{i+1}$ by:

$$c_i = \frac{2\left|(\vec{v}_{i+1} - \vec{v}_i) \times (\vec{v}_i - \vec{v}_{i-1})\right|}{\left|\vec{v}_{i+1} - \vec{v}_i\right|\left|\vec{v}_i - \vec{v}_{i-1}\right|\left|\vec{v}_{i+1} - \vec{v}_{i-1}\right|}$$

**Manual Validation**

Quantification of the same WT and *git-1* image dataset (n=10 WT and n=10 *git-1*) was performed by manual methods [5]. As an approximation to morphological classes of tips (Fig. 6E), quaternary branches and ectopic branches were quantified and normalized per 100µm of the primary branch length. Ectopic branches are defined as non-idealized terminal branches of menorahs. The number of ectopic branches of orders 2-4 (roughly corresponding to morphological classes 1-3), normalized per 100µm of the primary branch length in wild-type animals is 3.8±0.3 , while in *git-1(ok1848)* this number is significantly increased:6.6±0.4 ($p$<0.0001).. The number of quaternary terminal branches (roughly corresponding to morphological class 4) normalized per 100µm of the primary branch length is 18.1±0.6, while in *git-1(ok1848)* it is significantly decreased to 15.2±0.5 ($p$=0.0039)..

**Quantification of morphological changes in the PVD during development**

At the L4 stage the PVD shows its full stereotypic Candelabra-like pattern that is maintained throughout early adulthood [5,6]. Manual (non-computational) analysis of

L4 and 5-10-day-old worms demonstrated that PVD structure undergoes dynamic changes during aging including increased number of branches, appearance of high-order ectopic branches and disorganized structure of PVD candelabras [7]. Hence, we hypothesized that PVD would display structural changes already at L4 to young-adult transition. However, there is no available quantitative data comparing the PVD structure at these stages. In addition, since a single developmental stage transition might consist of minor changes in PVD structure, we assumed that computational analysis could detect and quantify changes that are otherwise missed by manual analyses of PVD images. We imaged anesthetized L4 and young adult worms and analyzed them using our software.

The software identified and classified different classes of PVD branches in L4 and young adult worms (Fig. S7A and B, respectively). As expected, young adult worms had increased neuronal length, number of junctions and number of tips (Fig. S7C-E and F-G). However, junction and tip density remained unchanged. (Figures S7H-I). These results suggest that during L4-young adult transition, the PVD scales up its number of junctions and tips proportionally to its length, such that vertex density is maintained. Scaling is a known phenomenon where a neuron expands its surface area and increases proportionally with the organism’s growth [8]. Finally, the contribution of specific morphological classes was examined. While all classes seem to contribute to the scaling up of PVD length and vertex number during the transition from L4 to young adult, some classes contribute more than others. Specifically, classes 1-3 contribute the most to the increase in length, class 1 contributes the most to the increase in number of junctions, and class 4 contributes the most to the increase in number of tips (Fig. S7E,J,K).

**Additional strain information**

All *git-1* mutant genotypes were confirmed by PCR; for genotyping *tm1962* the following primers were utilized: F1 5'-aaaactaatgattcgagagcagcga-3'; F2 5'-agaaagactttcaagtaccgaaactatg-3'; R 5'- ggagtctctacaatagccaatggaga-3' (WT products are 834 and 250 base pairs (bp), *tm1962* allele yields a single 373 bp band). A fragment of RB1540 *git-1(ok1848)* was sequenced by us using F 5'- aaaactaatgattcgagagcagcga-3' with 'outer right' primer of strain RB1540 R 5'- gttcccctatcatgccaaaa-3' to confirm an 891 bp deletion covering exons 12, 13 and 14. For genotyping *ok1848*, utilized F 5'-cacgcgccacattaaaaccc-3' and R 5'-acacgttccgaacctttcac-3' to amplify a 995 bp fragment in WT but not mutant animals.

*git-1(tm1962)* X was crossed into BP1021 [*him-5(e1490)* V; hmnIs133] and then *him-5(e1490)* was removed by crossing 2x into BP709. This strain was then outcrossed with N2 yielding BP1077 (*git-1(tm1962)* X; hmnIs133), and non-mutant worm siblings (1x outcrossed hmnIs133) that were confirmed by PCR and used as the 10 wildtype worms analyzed by the software. Harsh touch experiments utilize N2 or BP709 as WT, with additional controls as the harsh touch non-responsive CB1338 (*mec-3(e1338)*) and BP925 (*mec-4(e1611)* X, hmnIs133).

Crawling gait experiments utilize BP709 as WT, *git-1(tm1962)*, and RB1540 or BP1054 for *git-1(ok1848)*.

| | | Parameter | Value | Unit |
|---|---|---|---|---|
| Neuron Tracing | Convolution | Rotation range | [-70,70] | o |
| | | Rotation step | 5 | o |
| | | Minimum peak distance | 15 | o |
| | | Minimum peak prominence | 0.4 | |
| | | Forward increment length | 1 | pixels |
| | | Normalization minimum peak height | 0.07 | |
| | | Normalization minimum peak distance | 30 | o |
| | | Smoothing parameter | 0.05 | |
| | Rectangle size | Rectangle length-width ratio | 2 | |
| | | Rectangle width smoothing parameter | 0.5 | |
| | | Rectangle width sliding window | 6 | steps |
| | | Rectangle width scanning resolution | 0.035 | µm |
| | | Maximum distance from skeleton segment | 1.5 | µm |
| Preprocessing | CNN | Input_size | 64x64 | pixels |
| | | Dataset size (number of input samples) | 15,000 | |
| | | Number of source PVD images | 3 | |
| | | Number of input samples per image | 5000 | |
| | | Training set ratio | 0.8 | |
| | | Solver | Adam | |
| | | Maximum number of epochs | 100 | |
| | | Mini batch Size | 128 | |
| | | Initial learning rate | 0.001 | |
| | | Learning rate drop factor | 0.9 | |
| | | Learning rate drop period | 5 | Epochs |
| | | Shuffle | once | |
| | | Filter size | [3,3] | |
| | | Number of convolution layers | 2 | |
| | | Depth | 3 | |
| | | L2 Regularization | 0.0005 | |
| | Vertex Convolution | Rectangle length | 1.8 | µm |
| | | Circumference scanning resolution | 1 | o |
| | | Minimum peak distance | 20 | o |
| | | Minimum peak width | 5 | o |
| | | Minimum peak prominence | 0.15 | |
| | | Smoothing parameter | 0.99 | |
| Feature Extraction | Curvature | Smoothing parameter | 0.01 | |

Table S1. Parameters.

### Figure legends

**Table S1**. Parameter values of the algorithms.

**Figure S1**. Neural network architecture for semantic segmentation of images based on SegNet [3]. This architecture is made of an encoder subunit through which the input image shrinks in size, and a decoder subunit through which the image is upsampled back to its original size. The input layer gets 64x64 pixels patches of grayscale neuronal images and applies zero-centering normalization to them. Then, two convolution units (blue squares) are applied, with each unit consisting of a 2D 3x3 convolution layer, followed by a batch normalization layer (BN) and a rectified linear unit (ReLU) layer. This unit is followed by a 2x2 max pooling layer (green square) that reduces the size of the input by a factor of four. This sequence is repeated three times in the encoder, resulting in a 8x8 feature map with 64 features. The same sequence is repeated in the decoder, but with each max pooling layer replaced by an upsampling layer (purple square) that precedes each sequence of convolutions. Numbers below encoder layers show the output size of the pooling layer, and in the decoder they show the input to the upsampling layer. Finally, the softmax layer (yellow square) takes the output of the last ReLU layer and converts it into a probability distribution that sums up to 1. The last layer of the network is a pixel classification layer (red square). This layer computes the loss (cross-entropy) during training and performs the prediction of one of the predefined classes for new data.

**Figure S2.** The processing of a PVD image through the pipeline (Figure 1D).
**A** A maximum intensity, grayscale image of a PVD neuron. Arrows show the anterior (A), posterior (P), dorsal (D) and ventral (V) directions.
**B** The classification of the image into neuron and non-neuron pixels. CNN derived classification appears in red, manually added pixels in blue and manually removed pixels in yellow.
**C** Skeleton image derived from the binary image in B.
**D** The fully traced neuron.
**F** Segmentation of the traced neuron. Different segments appear in different colors.

**Figure S3**. Detection of the neuron midline and borderline used to define the PVD coordinate system.
**A** The neuron's trace is used to generate a single blob image. Then, the centerline and boundary of the blob are detected (dashed blue and green lines respectively). These are then refined using a sliding window along the approximated centerline, resulting in the final midline and boundary of the neuron (solid lines).
**B** A PVD color-coded for midline coordinate, from anterior (A) to posterior (P). Each neuron element is associating with a midline point by shortest distance, and colored according to its corresponding midline arclength from anterior to posterior.
**C** The distribution of midline orientation of junction rectangles. This distribution is different from the one of all neuron elements (Figure 3H), with peaks at 13.5° and 90°.

**Figure S4**. Algorithmically-derived morphological classes. Classes color-coded as in Fig. 4A-E: Class 1 (red), class 2 (green), class 3 (blue) and class 4 (yellow).
**A** Magnified regions of wild-type PVDs algorithmically classified into morphological classes. Arrows show classifications that do not match the conventional manual classification into Menorah orders.
**B-J** Visualization of the algorithmically-derived classification in nine PVD images of wild-type *C. elegans* worms (in addition to the one in Figure 4C).

**Figure S5**. The geometry of PVD junctions.
**A** Examples of PVD junction morphologies, showing a two-times larger neighborhood compared with Figure 5A, to demonstrate that process orientation often does not match junctional angles. Colors correspond to relative angle size: smallest (red), mid-size (green) and largest (blue).
**B** The variability in junction geometries is characterized by angular noise, determined from a Monte-Carlo simulation. Fit with simulated distributions gives a junction variability of 19° around the symmetrical configuration (120°-120°-120°), as indicated by the star symbol.

**Figure S6**. Algorithmically-derived morphological classes for *git-1(ok1848)* mutants.
**A-I** Visualization of the algorithmically-derived classification in nine PVD images of *git-1(ok1848) mutants* (in addition to the one in Figure 6A).

**Figure S7**. Quantification of morphological changes in the PVD during development.

**A** An image of a full PVD of an L4 worm, superimposed with color-coded morphological classes: class 1 (red), class 2 (green), class 3 (blue), class 4 (yellow).

**B** An image of a full PVD of a young-adult worm, superimposed with color-coded morphological classes, as in A.

**C** Distribution of neuronal length along the midline (head = 0), averaged across worms, for L4 (blue, n=5) and young-adult (red, n=5) worms.

**D** The total PVD length for L4 (blue) and young-adult (red) worms.

**E** The total PVD length for each morphological class for L4 (blue) and young-adult (red) worms.

**F** The total number of dendritic junctions, as described in D.

**G** The total number of dendritic tips, as described in D.

**H** The density of dendritic junctions, as described in D.

**I** The density of dendritic tips, as described in D.

**J** The total number of dendritic junctions for each morphological, as described in E.

**K** The total number of dendritic tips for each morphological, as described in E.

In D-K, statistics were calculated using the nonparametric Mann–Whitney test. $*p < 0.05$. n=5 L4 animals, with 1218 junctions and 1085 tips. n=5 young-adult animals, with 1610 junctions and 1410 tips. Bars show the mean value and error bars show the standard deviation.
