## Supplementary figures and images for "Neuron tracing and quantitative analyses of dendritic architecture reveal symmetrical three-way-junctions and phenotypes of *git-1* in *C. elegans*"

### SI figures merged

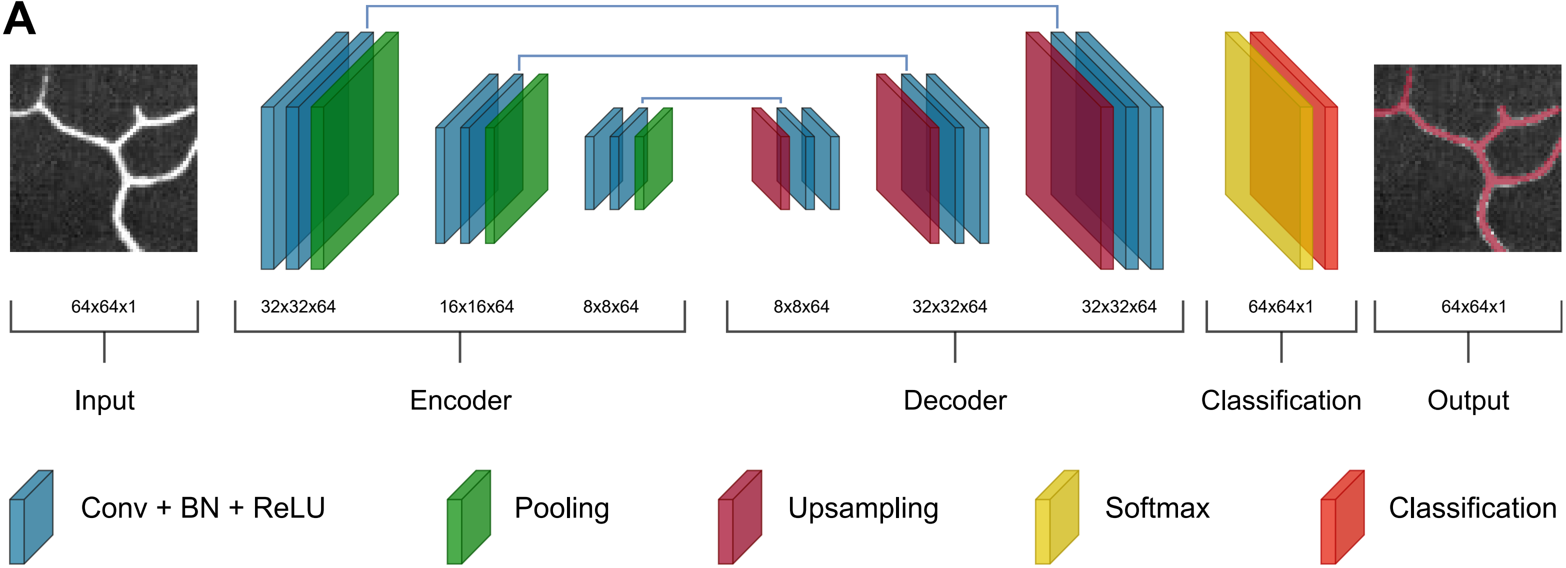

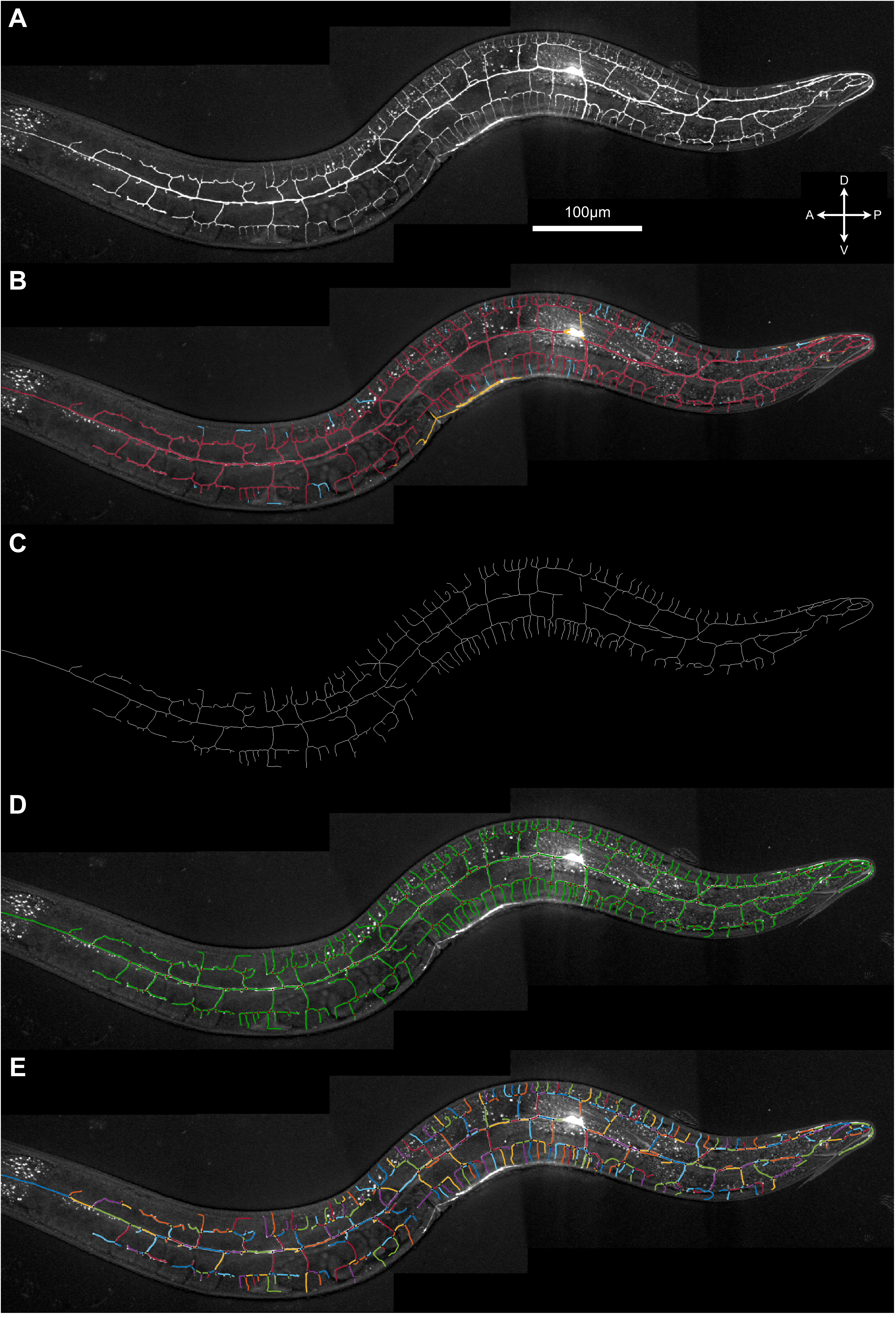

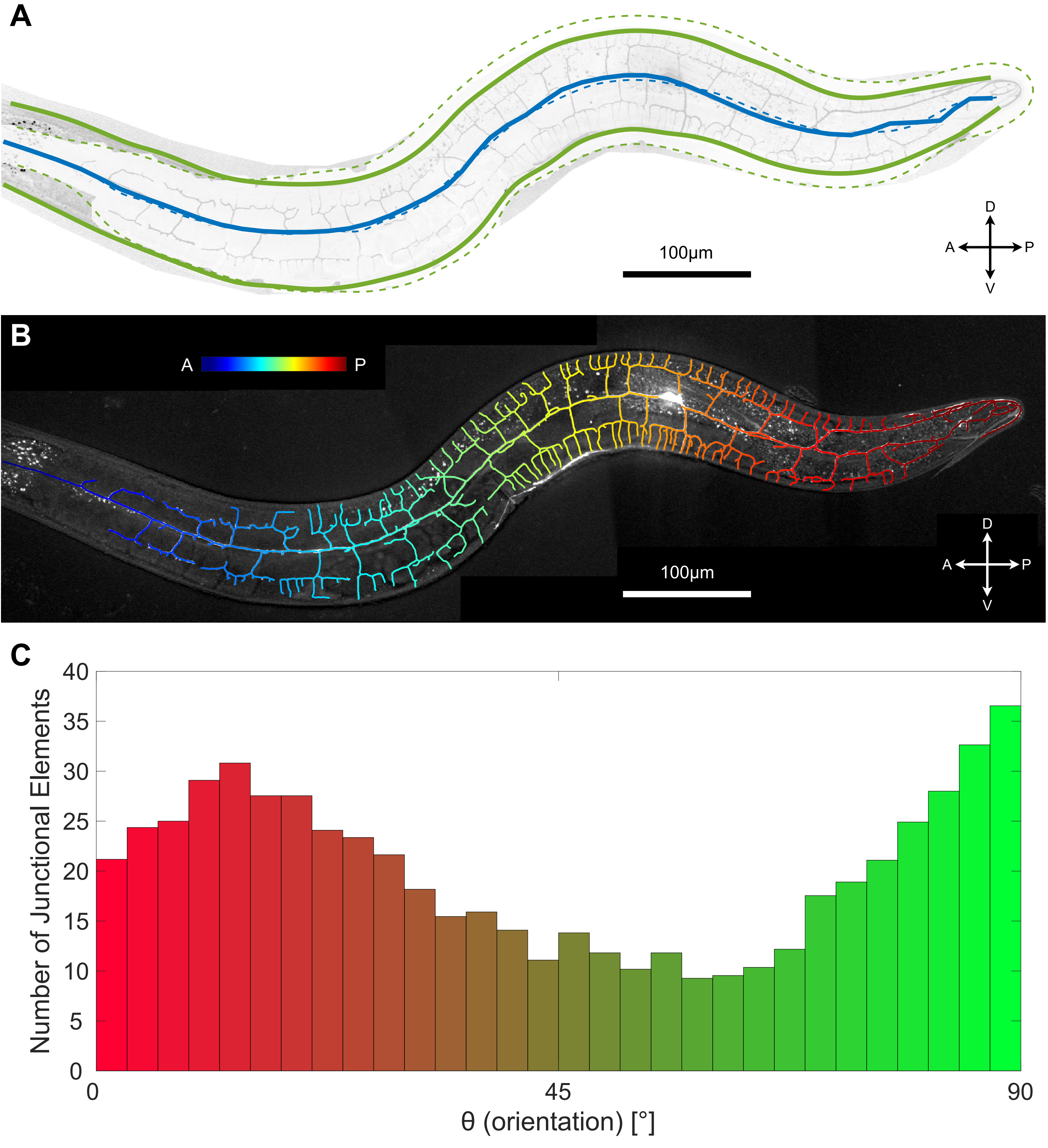

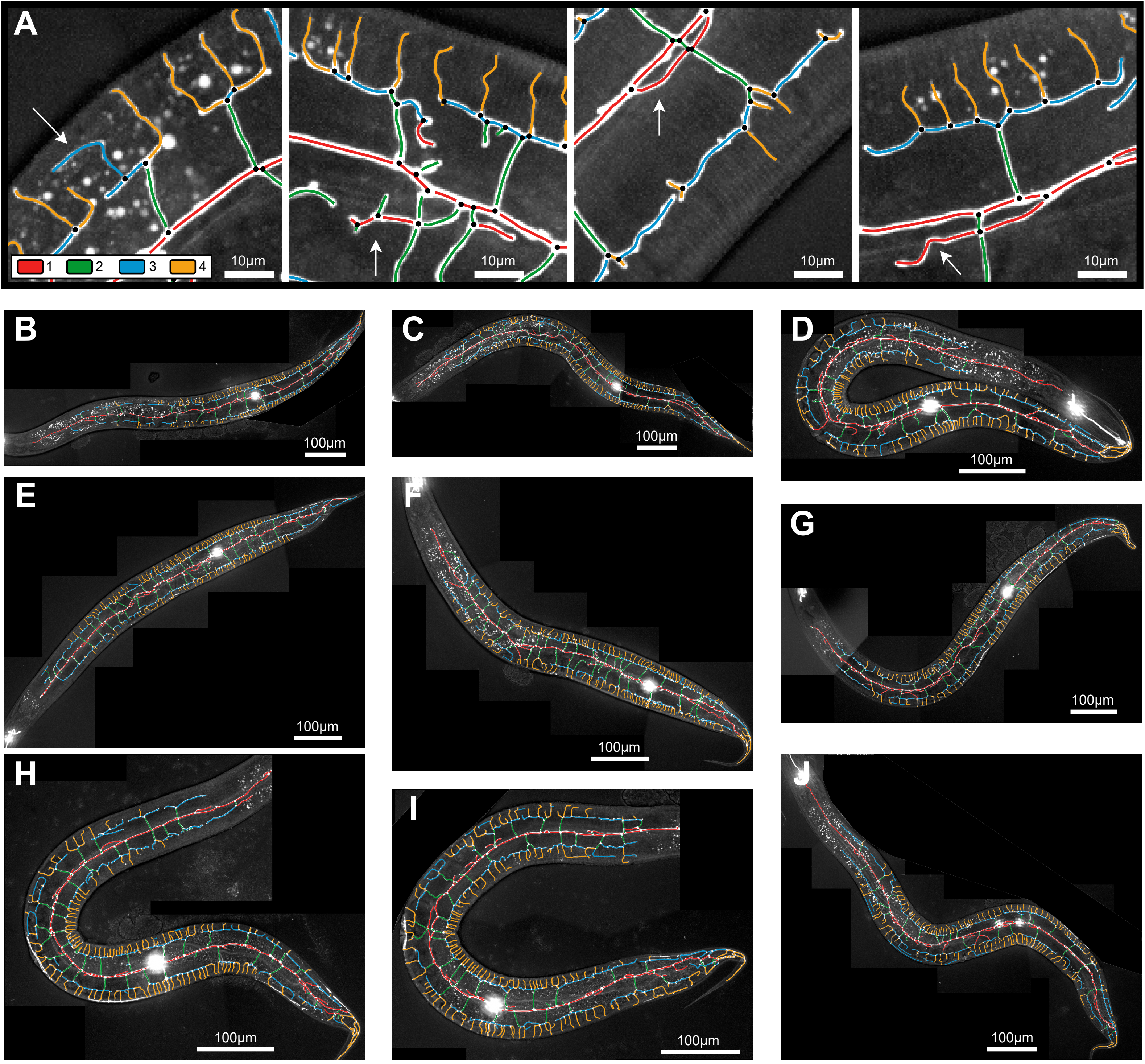

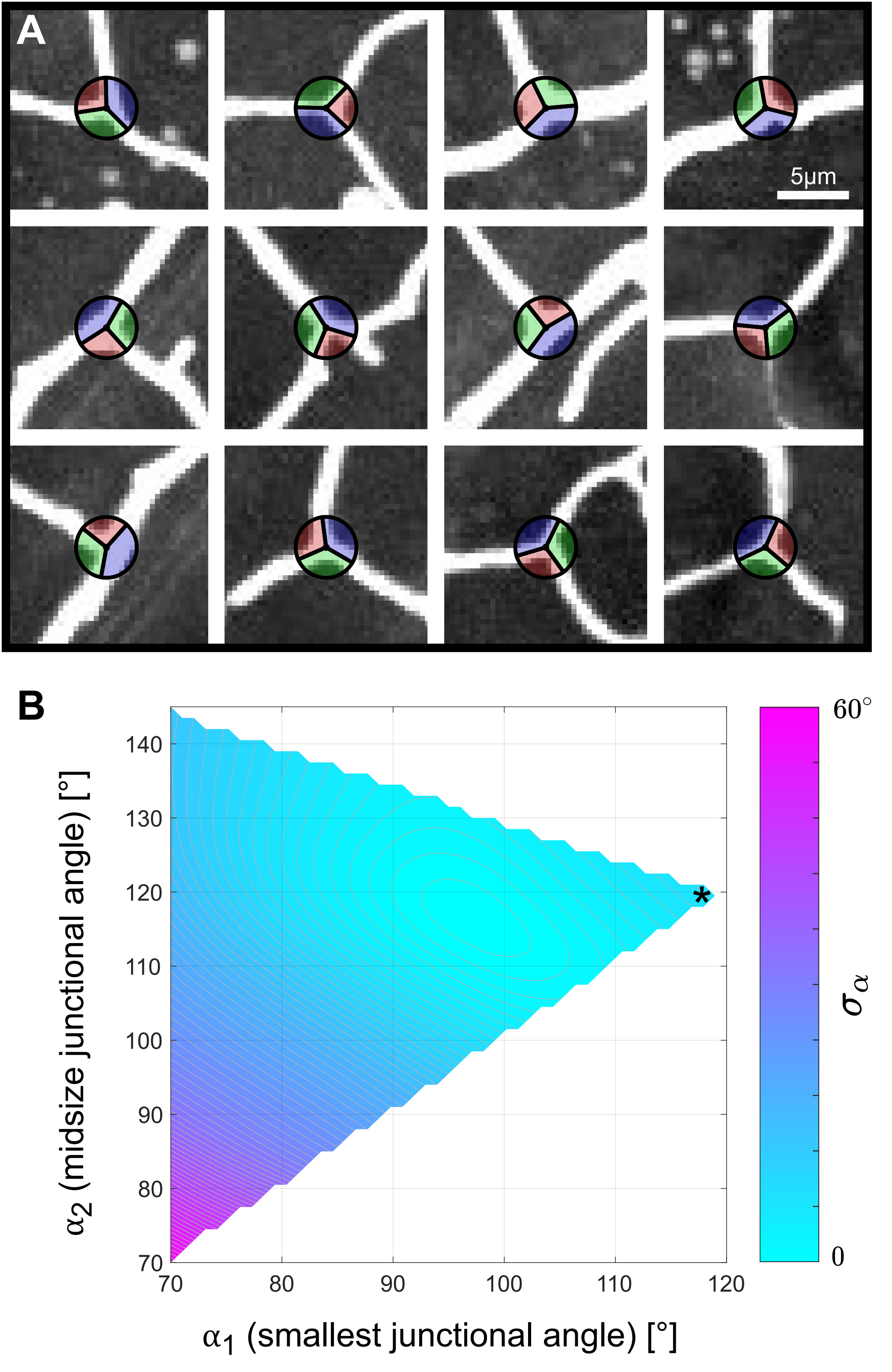

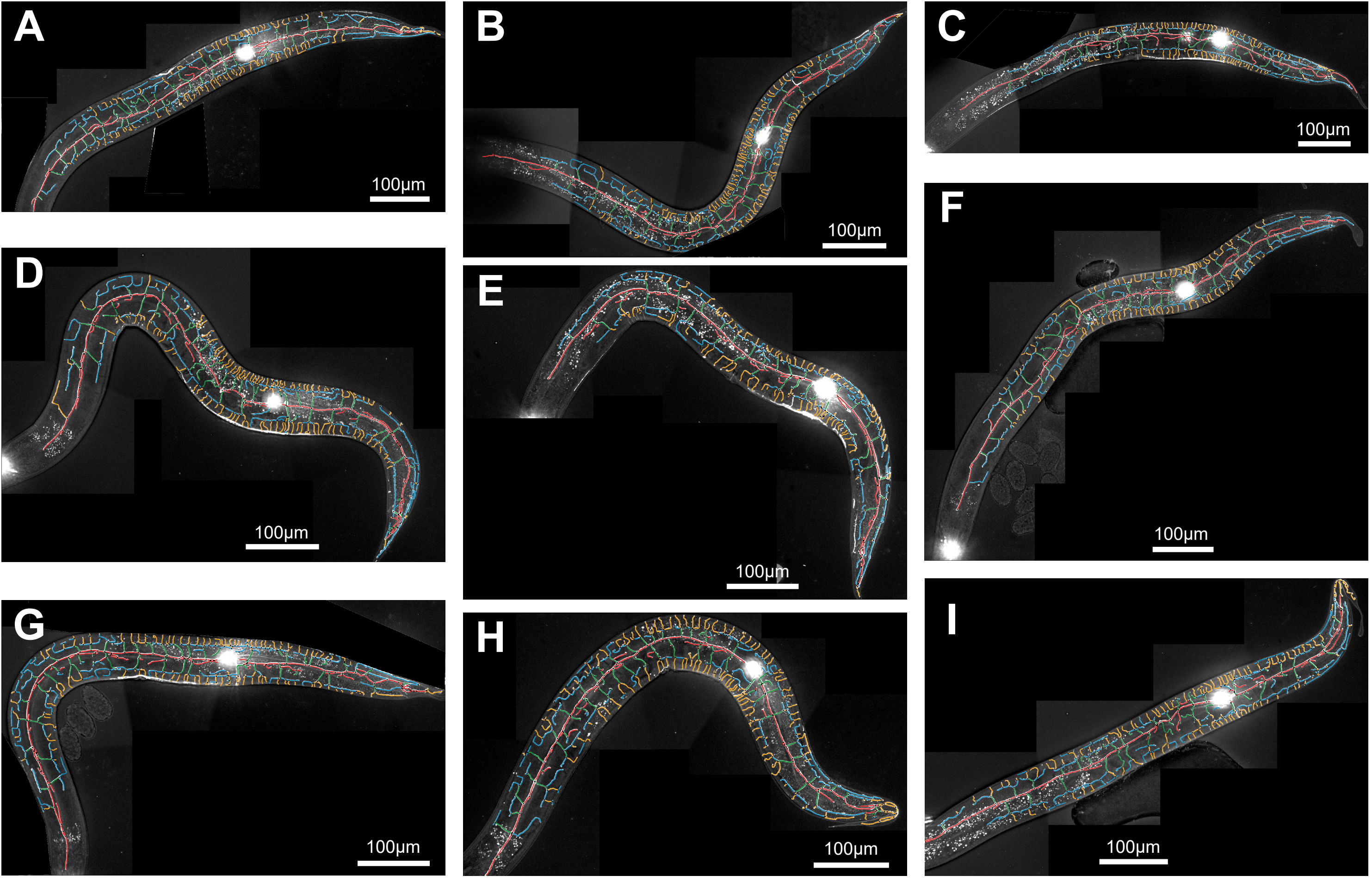

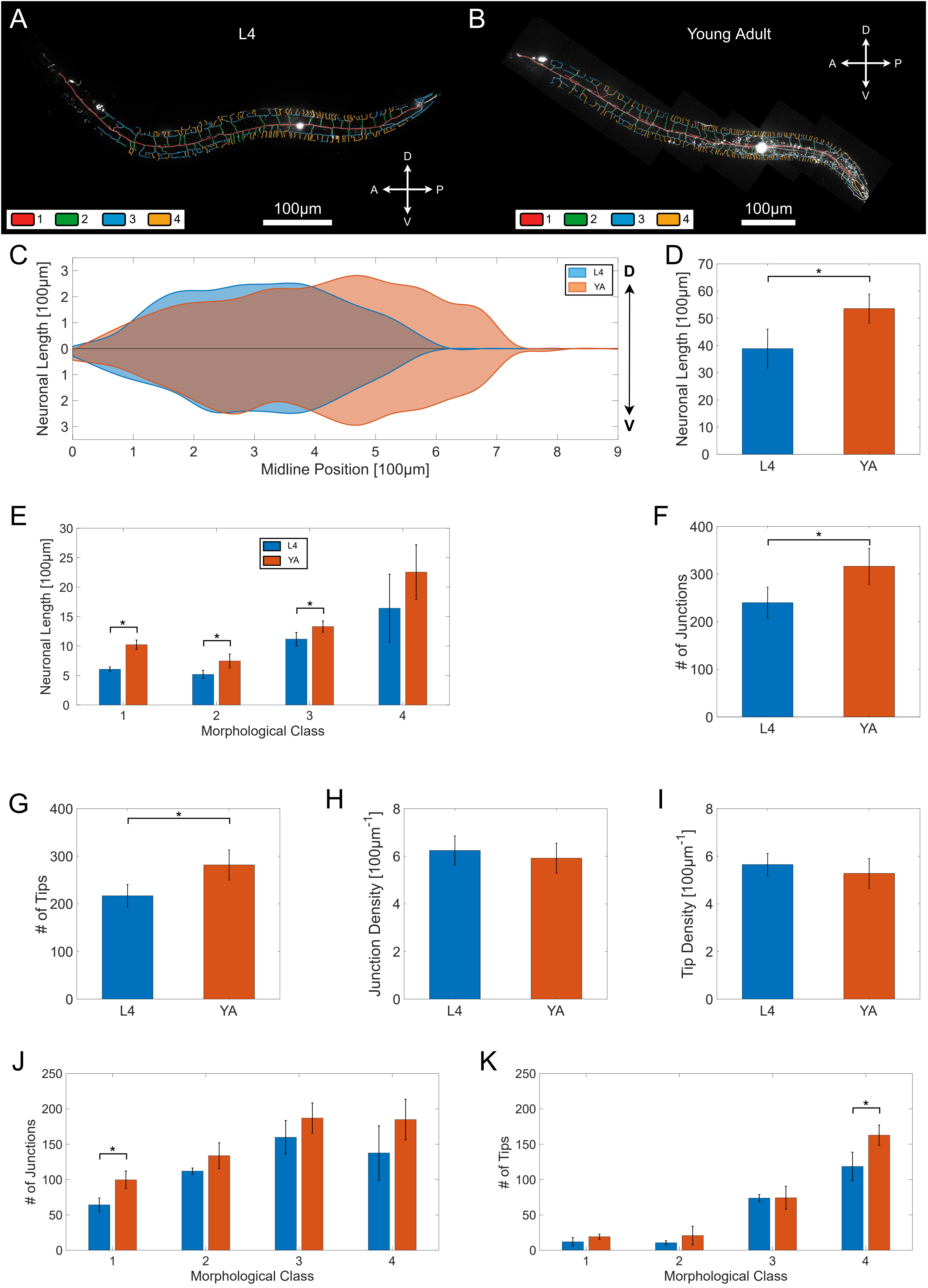
